# Seeing and smelling mates: multimodal integration and visual gating of chemical cues in female mate-location behavior in the prawn *Macrobrachium rosenbergii*

**DOI:** 10.64898/2026.05.12.723903

**Authors:** Felipe P. da Costa, Maria F. Arruda, Karina Ribeiro, Daniel M. A. Pessoa

## Abstract

Multimodal communication plays a central role in animal behavior, particularly when individuals must integrate information from different sensory channels to make rapid decisions. In aquatic environments, chemical and visual cues differ markedly in their spatial and temporal properties, such that chemical signals may be constrained by limited spatial resolution and temporal instability, potentially requiring visual information to reliably guide social decisions. In decapod crustaceans, both cue types are known to mediate reproduction, yet their relative contribution to mate-location behavior remains unclear. Here, we tested how visual and chemical cues from males influence mate-location behavior in females of the prawn *Macrobrachium rosenbergii*. Females were placed in a central arena and exposed to four stimulus configurations combining visual cues (a life-size photograph of a male or a control background) and chemical cues (water from an aquarium with or without a male). Attraction was quantified as the time spent in each half of the arena. Females showed no directional preference when exposed to chemical cues alone or when visual and chemical cues were spatially incongruent. In contrast, females spent significantly more time near male-associated stimuli only when visual and chemical cues were spatially congruent. These results indicate that mate-location behavior in this species depends on multimodal integration with a strong contextual dependence on visual information, which appears to gate the effectiveness of chemical cues. Spatially congruent multimodal signals are therefore necessary to guide orientation during mate search, suggesting that disruption of visual or chemical information in aquaculture systems may impair mating efficiency.

## INTRODUCTION

*Macrobrachium rosenbergii* is a catadromous prawn of great commercial importance (New, 2002; New et al., 2010), whose life cycle spans marine and freshwater environments, with larvae developing in estuarine and coastal waters before recruiting to freshwater habitats as juveniles (Ling, 1969; Upadhyay et al., 2014). This ontogenetic shift exposes individuals to contrasting ecological and sensory conditions that may shape their biology (Costa et al., 2023; Nilson et al., 1986). Correa and Thiel (2003) classified *M. rosenbergii* mating system as dominance neighborhoods, in which males compete locally for dominance, increasing their chances of future mating. Within this system, large blue claw males typically monopolize access to receptive females, whereas small males must adopt alternative strategies, such as actively searching for fertile females while avoiding direct confrontations with dominant individuals.

Fertilization in *M. rosenbergii* is restricted to a brief period following the female molt (Chow et al., 1982; Ling, 1969; Rao, 1965), imposing strong selective pressure on females to locate mates rapidly. Accordingly, females increase locomotor activity during premolt when mature eggs are present in the ovary (Peebles, 1979) and show attraction toward males (Bauer, 2011), and in some cases toward depressions in the substrate constructed by them (Smith & Sandifer, 1979). Female attraction is likely mediated by visual cues (Costa et al., 2025) and by the continuous or intermittent release of pheromones by males (Bauer, 2011), followed by the use of contact pheromones during close-range interactions (Bauer, 2011).

Upon detecting a receptive female, blue claw males approach with the body raised and antennae and claws extended forward (Chow et al., 1982; Ling, 1969). The male grasps the female between his claws (Chow et al., 1982; Ling, 1969; Rao, 1965), cleans her ventral surface (Ling, 1969; Rao, 1965), then turns her upside down to transfer the spermatophore (Chow et al., 1982; Ling, 1969; Rao, 1965). Mating and the presence of a spermatophore in the thelycum accelerate spawning, after which females carry the eggs in the brood chamber (Kruangkum et al., 2015). The presence of developing embryos further stimulates continued egg carrying until hatching (Kruangkum et al., 2015).

Chemical communication also plays a critical role in male reproductive physiology. The presence of premolt and postmolt females increases the expression of the insulin-like androgenic gland hormone (IAG) in the androgenic gland and neuropeptides associated with olfactory functions in male nervous tissues, promoting testicular cell proliferation, and these changes appear to be triggered by chemical substances released by females and detected via the short lateral antennules (Kruangkum et al., 2019; Kruangkum et al., 2025).

However, despite their central role in animal communication (Hay, 2009), chemical signals in aquatic environments can be temporally unreliable, as they may degrade rapidly after release and lose their behavioural relevance within minutes depending on environmental conditions such as solar radiation, temperature, and water chemistry (Chivers et al., 2013). Visual communication is also constrained, as turbidity can scatter light and reduce contrast, thereby limiting signal detection and transmission (Utne-Palm, 2002). In this context, multimodal signalling provides a useful framework for examining how visual and chemical cues are integrated, as signals may convey either redundant or complementary information (Hebets et al., 2016). When conveying similar information, cues may interact through equivalence, enhancement, or antagonism, whereas signals conveying different information may interact through independence, dominance, modulation, or emergence (Munoz & Blumstein, 2012; Partan & Marler, 1999). Despite this, the relative contribution of visual and chemical cues to male–female interactions in *M. rosenbergii* remains unknown.

To address this gap, we experimentally tested how females use visual and chemical information during mate-location behaviour by quantifying their time allocation within an experimental arena. Given the inherent constraints of chemical signalling, particularly its temporal instability and limited spatial resolution, we hypothesized that chemical cues would only be effective in guiding mate-location behaviour when accompanied by visual information, following predictions from multimodal signaling frameworks (Hebets et al., 2016; Munoz & Blumstein, 2012; Partan & Marler, 1999). Specifically, we predicted that: (1) females would not show an increased residence time in zones associated with male stimuli when only chemical cues were available; (2) females would not prefer a single zone when visual and chemical cues were presented on opposite sides of the aquarium; and (3) females would spend significantly more time in the zone associated with male stimuli when visual and chemical cues were presented together on the same side of the aquarium. Support for these predictions would indicate a modulation effect arising from multimodal cue integration (Munoz & Blumstein, 2012; Partan & Marler, 1999), rather than reliance on a single sensory modality, likely reflecting an adaptation shaped by natural selection to overcome constraints on signal transmission and ensure timely mate identification.

## MATERIAL AND METHODS

### Animal maintenance

Juvenile prawns were obtained from the Agricultural School of Jundiai, Rio Grande do Norte, Brazil, and transported to the Laboratory of Sensory Ecology of the Federal University of Rio Grande do Norte, where they were kept until they reached adulthood, as explained in Costa et al. (2025). Animals were housed in collective aquaria (1.50 × 0.80 m) containing a sand substrate, continuous aeration, and a filtration system composed of sponge, activated carbon, and ceramic rings. After the animals had fully developed, a partition system made of PVC pipes, plastic mesh, and mosquito netting was installed to prevent aggressive interactions, allowing chemical and visual contact but preventing physical interactions. Each prawn was provided with an individual plastic shelter. We identified the sex of the prawns by the presence of appendix masculina, if a male, or ovary, if a female.

Aquaria were maintained under an approximately 12 h light : 12 h dark photoperiod, as recommended by Wei et al. (2025). Prawns were fed twice daily with commercial feed. Water quality parameters were maintained within levels suitable for the species (ammonia: 0 ppm; temperature: 26–27 °C; pH: 7.0–7.2).

### Experimental procedure

To test the hypothesis that effective mate location by females requires the simultaneous availability of visual and chemical cues, we quantified female time allocation in response to different combinations of male-derived stimuli. Trials were conducted in a central aquarium (0.50 × 0.30 m) without substrate, flanked by two adjacent aquaria (0.20 × 0.35 m), also without substrate. Adjacent aquaria contained either a blue claw male or clean water that had never been in contact with prawns.

Chemical cues were operationalized as water flowing from the adjacent aquaria into the central aquarium through a siphon system. Visual cues were operationalized as a life-size photograph (rostrum–caudal length: 13.0 cm) of the same blue claw male used as the chemical cue source, displayed against a black background, or as the black background alone. A single male was used throughout the experiment to control for variation in male phenotype (Bushmann & Atema, 2000).

To test each operational prediction, four stimulus configurations were used (Fig. 1): (1) absence of visual and chemical cues on both sides (control); (2) chemical cues on one side only; (3) visual and chemical cues presented together on one side; and (4) chemical cues on one side and visual cues on the opposite side. These configurations directly corresponded to predictions that females would not show a directional bias when exposed to unimodal chemical cues or spatially separated chemical and visual cues, but would show a directional bias when cues were combined. Importantly, previous work has demonstrated that females can use visual cues in isolation for mate detection, indicating that visual information alone can be sufficient to guide orientation in this species under certain conditions (Costa et al., 2025).

**Figure 1.**
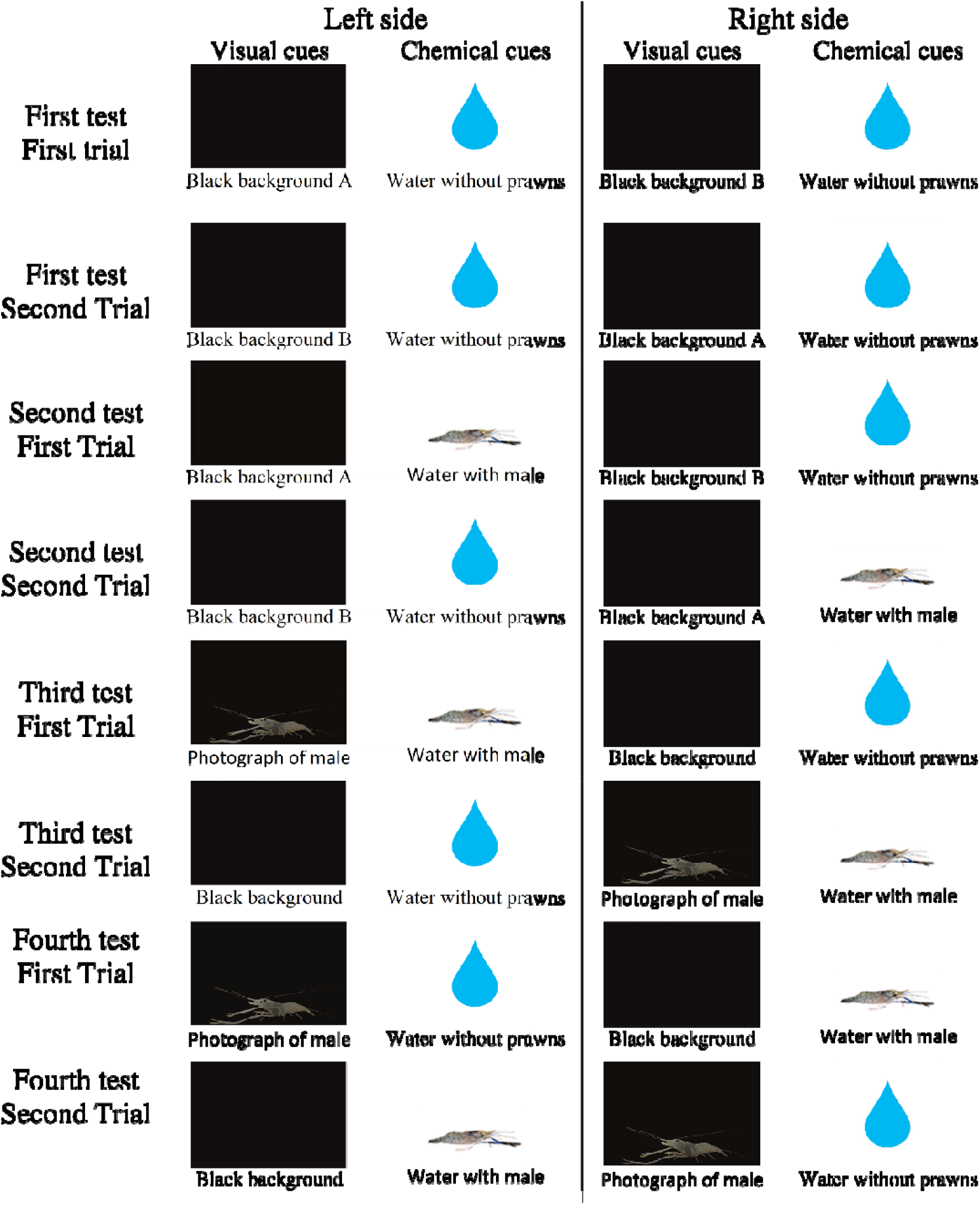
Stimulus combinations used in the tests of attraction to visual and chemical cues from males. Four experimental configurations were tested: (1) control condition, in which both sides of the central aquarium contained no visual or chemical male cues (black background and water without prawns); (2) chemical cue condition, in which male-conditioned water was presented on one side and control water on the other; (3) combined-cue condition, in which a male photograph and male-conditioned water were presented together on one side, with control stimuli on the opposite side; and (4) spatially incongruent condition, in which male-conditioned water was presented on one side and the male photograph was presented on the opposite side, both contrasted against control stimuli. In all trials, the side (left or right) of each stimulus combination was alternated between replicates to avoid side bias.

Females in ovarian development stage V (Martins et al., 2007) were placed individually in the central aquarium, and the male (when used) was placed in one of the adjacent aquaria, followed by a 30-min acclimation period. After acclimation, visual stimuli were positioned between aquaria and, simultaneously, chemical cues were initiated. The central aquarium was not aerated to prevent interference with chemical cue dispersion.

Female behavior was recorded from above for 13 min. To operationalize attraction, we quantified the total time spent by females in each half of the central aquarium. Analyses were conducted for the final 10 min of each trial and, separately, for the last 3 min, when chemical cue concentration was expected to be highest. Time spent straddling the midline was excluded. A greater proportion of time spent in one half of the aquarium was interpreted as evidence of directional orientation toward the corresponding stimulus combination.

At the start of each trial, the central aquarium contained 13.5 L of water, and each adjacent aquarium contained 17.5 L. After 13 min, 3.5 L from each adjacent aquarium had flowed into the central aquarium. After each trial, animals were returned to their housing aquaria, and all experimental aquaria were dried and refilled with fresh water to prevent cue carryover.

Nine females were tested, each undergoing eight trials over three consecutive days (maximum of three trials per day), with at least 2 h between trials. The side of stimulus presentation (left or right) was alternated between replicate trials to control for side biases.

### Statistical analysis

To test each operational prediction, we used generalized estimating equations (GEE) with a gamma distribution and log link function to compare the time females spent in each half of the aquarium within each stimulus configuration. The gamma distribution was selected because the response variable was continuous and positively skewed and yielded lower Quasi-likelihood under Independence Model Criterion (QIC) values than linear models. To eliminate zero values and improve model fit, 100 s were added to each observation. Statistical significance was set at α = 0.05. Analyses were conducted using SPSS 25.

## RESULTS

When female attraction was evaluated over the 10-min observation period, no significant differences were detected between stimulus in any experimental condition (all p ≥ 0.140).

In contrast, analyses restricted to the final 3 min of each trial, when females had received a greater volume of water from the adjacent aquaria, revealed a significant effect of stimulus configuration. Females spent significantly more time in the half of the aquarium containing both visual and chemical cues from the male (111.11 ± 11.19 s) than in the half containing a black background and water without prawns (66.28 ± 11.26 s; Fig. 2) (Wald = 3.931; df = 1; p = 0.047).

**Figure 2.**
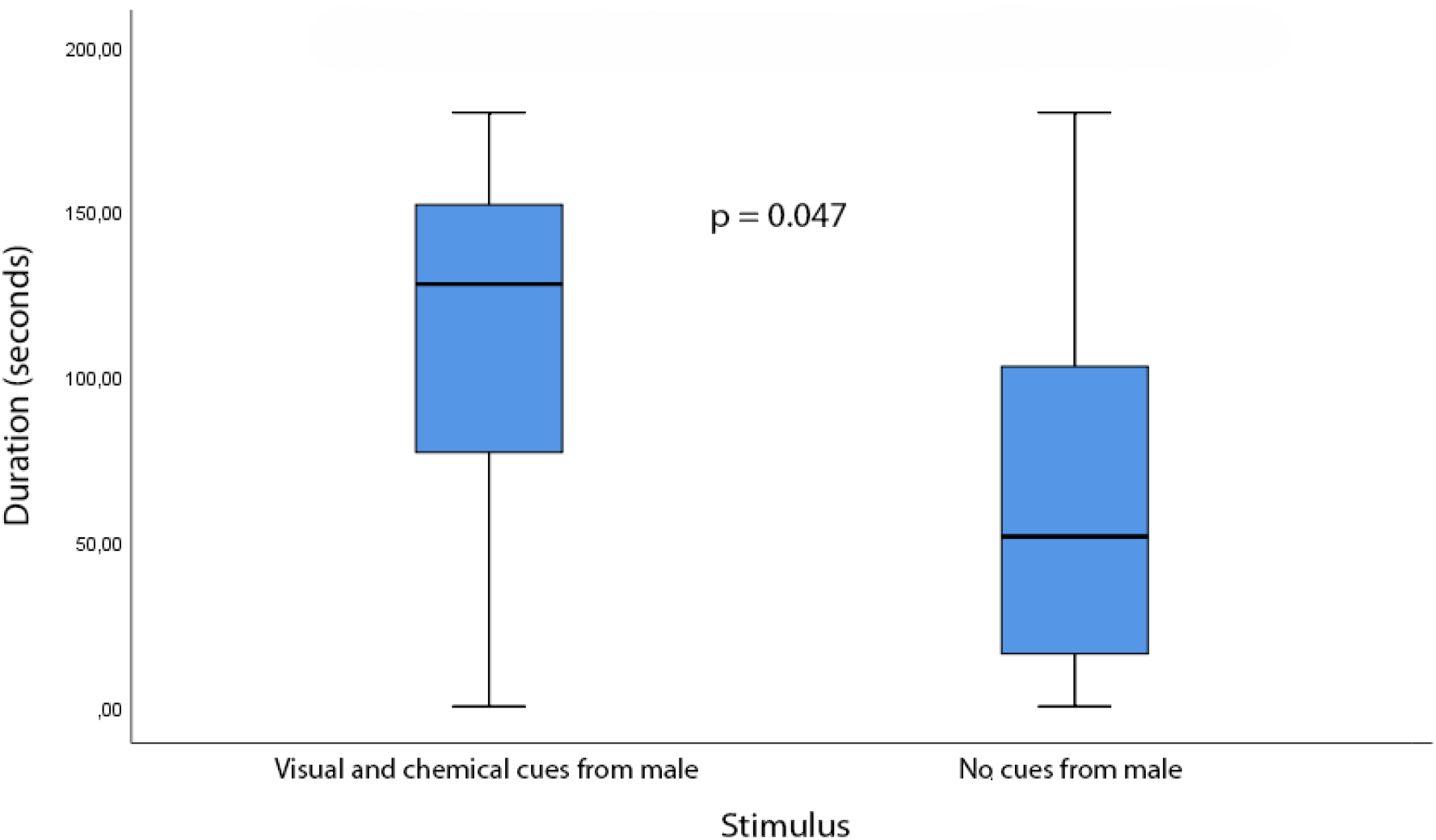
Female attraction to combined visual and chemical cues from a male. Boxplots show the total time females spent in the half of the central aquarium presenting a photograph of a male and receiving water from an adjacent aquarium containing a live male, and in the opposite half presenting a black background and receiving water from an aquarium without prawns. Horizontal lines within boxes indicate medians; box limits represent first and third quartiles; whiskers indicate minimum and maximum values.

No significant preference was observed in any of the other stimulus configurations, including those presenting chemical cues alone or visual and chemical cues on opposite sides of the aquarium (all p ≥ 0.081).

## DISCUSSION

Our results show that mate location by female *Macrobrachium rosenbergii* emerges from a behaviorally asymmetric process of multisensory integration, in which chemical information alone is insufficient and requires concurrent visual input to guide orientation, consistent with a hierarchical multimodal system characterized by visual dominance and context-dependent modulation by chemical cues (Munoz & Blumstein, 2012; Partan & Marler, 1999). Females exhibited attraction only when visual and chemical cues from males were spatially congruent, whereas isolated chemical cues or spatially incongruent combinations failed to elicit a directional response, despite previous evidence that visual cues alone can support mate detection in this species (Costa et al., 2025). This pattern indicates that visual information plays a primary role in structuring mate-search behavior, effectively gating the behavioral relevance of chemical signals under our experimental conditions. Together, these results support the hypothesis that mate-location is not driven by independent sensory modalities, but by an integrated and hierarchically organized perceptual process in which chemical cues contribute only when embedded within a coherent visual context.

This finding highlights that chemical cues alone are insufficient to support spatial decision-making under the experimental conditions used here. Although chemical signals can convey information about male presence or reproductive status, their directional reliability is often limited by turbulent flow and plume fragmentation (Atema, 2012). Visual cues, in turn, provide precise spatial information but may lack biological relevance when presented in isolation. The attraction observed only when both cues co-occurred suggests that females use chemical cues to confirm biological significance and visual cues to resolve spatial orientation, a process consistent with models of sensory weighting and cue validation in animal decision-making (Sheppard et al., 2013).

Our results differ from those reported by Al-mohsen (2009), who observed attraction of females to male chemical cues alone. This discrepancy likely reflects differences in how behavioral responses were quantified. Whereas our design required females to make a continuous spatial choice between two competing stimulus fields, Al-mohsen (2009) measured latency to reach a single water inlet. Under such conditions, chemical cues alone may increase general activity without providing sufficient directional information for side preference. This distinction emphasizes the importance of experimental context when interpreting behavioral responses to sensory cues and reinforces the view that attraction and orientation are separable behavioral processes.

Evidence from other crustaceans supports the importance of multimodal integration in mediating social interactions. In snapping shrimp (*Alpheus heterochaelis*), chemical cues alone fail to guide directional responses, whereas visual cues enable accurate orientation during social encounters (Hughes, 1996a, 1996b). Similarly, in the American lobster (*Homarus americanus*), effective social recognition depends on the combined use of visual and chemical information (Bruce et al., 2018). Together with our findings, these studies indicate that multimodal sensory integration is a common mechanism underlying spatial decision-making in decapod crustaceans.

The temporal dynamics observed in our study further reinforce this interpretation. Females expressed attraction to the combined cues only during the final 3 min of observation, when chemical cues were likely more concentrated. This suggests that chemical information modulates responsiveness to visual cues rather than directly determining orientation, consistent with a sequential decision process in which sensory inputs are integrated over time before a directional response is expressed.

From an applied perspective, these results have direct implications for the design of breeding systems for *M. rosenbergii*. Artificial environments that restrict either visual contact (e.g., low light levels, high turbidity) or chemical transmission (e.g., excessive filtration or unidirectional water flow) may disrupt the natural behavioral processes underlying mate location. Breeding facilities should therefore promote simultaneous visual exposure and localized chemical exchange between males and receptive females to facilitate efficient mate finding. Incorporating these principles into aquaculture systems may improve mating success and reduce the time females spend searching for mates during their brief receptive period. Future studies should explore how females integrate multimodal cues when choosing among males of different sizes or morphotypes and how environmental variables such as water flow and turbidity shape this decision-making process.

## FUNDING

This study was financed in part by the Coordenacao de Aperfeicoamento de Pessoal de Nivel Superior – Brazil (CAPES), Finance Codes 001, and 043/2012, that provided a Ph.D. Scholarship to F.P.C.; and by Conselho Nacional de Desenvolvimento Cientifico e Tecnologico – Brazil (CNPQ), Finance Codes 478222/2006-8, 25674/2009 and 474392/2013-9, that provided a Research Productivity Fellowship to D.M.A.P.

## ACKNOWLEDGEMENTS

We would like to thank Felipe N. Castro for his advice regarding the statistical analyses. We would also like to thank Ana L. F. Emerenciano, Kevin J. C. Martins, Lara M. B. Costa, Maria C. O. Guerreiro, Moisés O. Souza, and Whennita H. O. Andrade for their help with the care of the prawns. We would like to thank Coordenacao de Aperfeicoamento de Pessoal de Nivel Superior – Brazil (CAPES), Finance Codes 001, and 043/2012, that financed in part our study and that also provided a PhD. Scholarship to F.P.C. We would also like to thank Conselho Nacional de Desenvolvimento Cientifico e Tecnologico – Brazil (CNPQ), Finance Codes 478222/2006-8, 25674/2009 and 474392/2013-9, that financed in part our study and that also provided a Research Productivity Fellowship to D.M.A.P.

## DECLARATIONS

### Confliict of interest

none.

### Data availability statement

The data supporting this research will be available at Zenodo in case of acceptance of the manuscript.

### Ethical statement

We follow ARRIVE guidelines and respect Brazilian laws. Our research was approved by the ethics committee of our institution (protocol 042/2018).

### Declaration of generative AI and AI-assisted technologies in the manuscript preparation process

Generative artificial intelligence tools were used solely to assist with language editing and to improve clarity and readability of the manuscript. These tools were not used to generate, analyze, or interpret data, nor to develop hypotheses, experimental design, or conclusions. All scientific content and intellectual contributions are exclusively attributable to the authors, who take full responsibility for the integrity and accuracy of the work.

## Notes

### Competing Interest Statement

The authors have declared no competing interest.

## REFERENCES

Al-mohsen, I., 2009. *Macrobrachium rosenbergii* (de Man 1879): the antennal gland and the role of pheromones in mating behaviour. PhD Thesis. University of Stirling.

Atema, J., 2012. Aquatic odour dispersal fields: opportunities and limits of detection, communication, and navigation, in: Brönmark C, Hansson L-A (Eds.) Chemical Ecology in Aquatic Systems. Oxford University Press, New York, pp 1–18.

Bauer, R.T., 2011. Chemical communication in decapod shrimps: The influence of mating and social systems on the relative importance of olfactory and contact pheromones, in: Breithaupt, T., Thiel, M. (Eds.) Chemical Communication in Crustaceans. Springer, New York, pp 277–296.

Bruce, M., Doherty. T., Kaplan. J., Sutherland. C., Atema. J., 2018. American lobsters, *Homarus americanus*, use vision for initial opponent evaluation and subsequent memory. Bull. Mar. Sci. 94(3), 517–532. 10.5343/bms.2017.1147

Bushmann, P.J., Atema, J., 2000. Chemically mediated mate location and evaluation in the lobster, *Homarus americanus*. J. Chem. Ecol. 26, 883–899. 10.1023/A:1005404107918

Chivers, D.P., McCormick, M.I., Mitchell, M.D., Ramasamy, R.A., Ferrari, M.C.O., 2013. Temporal variation in the effectiveness of chemical alarm cues: the role of degradation in aquatic environments. Anim. Behav. 86, 1009–1015. 10.1016/j.anbehav.2013.08.014

Chow, S., Ogasawara, Y., Taki, Y. 1982. Male Reproductive System and Fertilization of the Palaemonid Shrimp *Macrobrachium rosenbergii*. Bull. Jpn Soc. Sci. Fish. 48(2), 177–183. 10.2331/suisan.48.177

Correa, C., Thiel, M. 2003. Mating systems in caridean shrimp (Decapoda: Caridea) and their evolutionary consequences for sexual dimorphism and reproductive biology. Rev. Chil. Hist. Nat. 76(2), 187–203. 10.4067/S0716-078×2003000200006

Costa, F.P., Arruda, M.F., Ribeiro, K., Pessoa, D.M.A., 2023. Influence of color and brightness on ontogenetic shelter preference by the prawn Macrobrachium rosenbergii (Decapoda: Palaemonidae). Zoologia (Curitiba) 40, e22023. 10.1590/S1984-4689.v40.e22023

Costa, F.P., Arruda, M.F., Ribeiro, K., Pessoa, D.M.A., 2025. The importance of color and body size for reproductive decision making by males and females of the giant river prawn, *Macrobrachium rosenbergii* (de Man, 1879) (Decapoda, Caridea, Palaemonidae). Behav. Process. 225, 105137. 10.1016/j.beproc.2024.105137

Hay, M.E., 2009. Marine chemical ecology: chemical signals and cues structure marine populations, communities, and ecosystems. Annu. Rev. Mar. Sci. 1, 193–212. 10.1146/annurev.marine.010908.163708

Hebets, E.A., Barron, A.B., Balakrishnan, C.N., Hauber, M.E., Mason, P.H., Hoke, K.L., 2016. A systems approach to animal communication. Proc. R. Soc. B Biol. Sci. 283(1826), 20152889. 10.1098/rspb.2015.2889

Hughes, M., 1996a. Size assessment via a visual signal in snapping shrimp. Behav. Ecol. Sociobiol. 38, 51–57. 10.1007/s002650050216

Hughes, M., 1996b. The function of concurrent signals: Visual and chemical communication in snapping shrimp. Anim. Behav. 52(2), 247–257. 10.1006/anbe.1996.0170

Kruangkum, T., Jaiboon, K., Vanichviriyakit, R., Sobhon, P., Chotwiwatthanakun, C., 2025. Upregulation of olfactory-related neuropeptide transcripts in male *Macrobrachium rosenbergii* in correlation to pheromone perception from molting females. Comp. Biochem. Physiol., Part A 302(2025), 111812. 10.1016/j.cbpa.2025.111812

Kruangkum, T., Saetan, J., Chotwiwatthanakun, C., Vanichviriyakit, R., Thongrod, S., Thintharua, P., Tulyananda, T., Sobhon, P., 2019. Co-culture of males with late premolt to early postmolt female giant freshwater prawns, *Macrobrachium rosenbergii* resulted in greater abundances of insulin-like androgenic gland hormone and gonad maturation in male prawns as a result of olfactory receptor. Anim. Reprod. Sci. 210, 106198. 10.1016/j.anireprosci.2019.106198

Kruangkum, T., Vanichviriyakit, R., Chotwiwatthanakun, C., Saetan, J., Tinikul, Y., Wanichanon, C., Cummins, S.F., Hanna, P.J., Sobhon, P., 2015. Spermatophore affects the egg-spawning and egg-carrying behavior in the female giant freshwater prawn, *Macrobrachium rosenbergii*. Anim. Reprod. Sci. 161, 129–137. 10.1016/j.anireprosci.2015.08.015

Ling, S.W., 1969. The general biology and development of *Macrobrachium rosenbergii*. FAO Fish. Rep. 57, 589–606.

Martins, J., Ribeiro, K., Rangel-Figueiredo, T., Coimbra, J., 2007. Reproductive cycle, ovarian development, and vertebrate-type steroids profile in the freshwater prawn *Macrobrachium rosenbergii*. J. Crustac. Biol. 27(2), 220–228. 10.1651/C-2597.1

Munoz, N.E., Blumstein, D.T., 2012. Multisensory perception in uncertain environments. Behav. Ecol. 23(3), 457–462. 10.1093/beheco/arr220

New, M.B., 2002. Farming freshwater prawns: a manual for the culture of the giant river prawn (Macrobrachium rosenbergii). FAO Fisheries Technical Paper 428. Rome. https://www.fao.org/3/y4100e/y4100e.pdf

New, M.B., Valenti, W.C., Tidwell, J.H., D’Abramo, L.R., Kutty, M.N., 2010. Freshwater prawns: biology and farmig. Wiley-Blackwell, Oxford.

Nilsson, D.E., Hallberg, E., Elofsson, R., 1986. The ontogenetic development of refracting superposition eyes in crustaceans: Transformation of optical design. Tissue Cell 18, 509–519. 10.1016/0040-8166(86)90017-0

Partan, S., Marler, P., 1999. Communication goes multimodal. Science 283(5406), 1272–1273. https://www.science.org/doi/10.1126/science.283.5406.1272

Peebles, J.B., 1979. Molting, Movement, and Dispersion in the Freshwater Prawn *Macrobrachium rosenbergii*. J. Fish. Res. Board Can. 36(9), 1080–1089. 10.1139/f79-151

Rao, R., 1965. Breeding behaviour in *Macrobrachium rosenbergii* (De Man). Fish. Tecnol. 2(1), 19–25.

Sheppard, J., Raposo, D., Churchland, A.K., 2013. Dynamic weighting of multisensory stimuli shapes decision making in rats and humans. J. of vision 13(6), 1–19. 10.1167/13.6.4

Smith, T.I.J., Sandifer, P.A., 1979. Breeding depressions in culture ponds for Malaysian prawns. Aquaculture 18(1), 51–57. 10.1016/0044-8486(79)90101-7

Upadhyay, A., Kulkarni, B.G., Pandey, A.K., 2014. Migration in prawns with special reference to light and water current as inducers in *Macrobrachium rosenbergii*. J. Exp. Zool. 17, 33–48.

Utne-Palm, A.C., 2002. Visual feeding of fish in a turbid environment: Physical and behavioural aspects. Mar. Freshw. Behav. Physiol. 35, 111–128. 10.1080/10236240290025644

Wei, J., Wang, Y., Hong, K., Huang, C., Nie, X., Liu, M., Zhou, Q., Su, Q., Mai, Z., Liufu, B., Li, H., Zhu, X., Yu, L., 2025. Effects of photoperiod on survival, growth and germ cell related gene expression of juvenile giant freshwater prawn, *Macrobrachium rosenbergii*. Invertebr. Reprod. Dev. 69(4) 1–10. 10.1080/07924259.2025.2594076

